# Utilising the boundary layer to help restore the connectivity of fish habitats and populations

**DOI:** 10.1101/332338

**Authors:** Jabin R. Watson, Harriet R. Goodrich, Rebecca L. Cramp, Matthew A. Gordos, Craig E. Franklin

## Abstract

**Significance:** Habitat fragmentation is a significant contributor to the worldwide decline of freshwater ecosystem health, the most pervasive cause of which is culverts. Culverts act as a barrier to fish movement, impacting feeding, predator avoidance, spawning, and community structures. Here we show that a common remediation strategy that involves baffles, is detrimental to the successful passage of small bodied and juvenile fish at high velocities. To remedy this widespread problem, we present a novel remediation design that benefits a range of small-bodied species and juvenile fish at the same high velocities, regardless of morphology or ecological niche. The application of this remediation design may be expanded to any smooth surfaced anthropogenic structure, to improve fish passage and restore ecosystem functionality.

**Abstract:** Culverts are a major cause of habitat fragmentation in freshwater ecosystems, are a barrier to fish movement, and are regarded as a significant contributor in the decline of freshwater fish populations globally. To try to address this, various culvert remediation designs have been implemented, including the installation of vertical baffles and the provision of naturalistic (rock) substrates. While remediation strategies generally aim to reduce the velocity of water flowing through the structure, there is often resistance to their use because the resultant reduction in culvert discharge can negatively impact upstream flooding while also resulting in debris clogging and increased culvert maintenance costs. In addition, baffles markedly increase water turbulence that may be detrimental to passage by some fish species or size classes. Here we present some novel remediation designs that exploit the reduced water velocity in boundary layers along the culvert wall to enhance fish passage without significantly compromising discharge capacity. These longitudinal designs produce an expanded reduced velocity zone along the culvert margins that generate minimal turbulence. We show that these novel designs are significantly advantageous to the swimming endurance and traversability for six small-bodied Australian fish species. We also provide data on how and why some culvert baffle designs may impede small-bodied fish passage. This data scales with increasing water velocity, encompassing inter-specific differences in swimming capacity. These results have broad implications for fish community structure and the requirement of juvenile cohort of large-bodied commercially important species where baffles have been implemented to facilitate fish passage.

## Introduction

Fish are a ubiquitous component of freshwater ecosystems, the diversity of which means we rely them directly for food, but also on their ecosystem services by maintaining system health, functionality and robustness (Gordon et al. 2018, Obregón et al. 2018, Rodríguez-Lozano et al. 2015). Despite this, freshwater systems are one of the most threatened by human activities (Carpenter et al. 2011). Regulating the flow of freshwater systems to control water access has severely fragmented fish habitats worldwide and threatens the persistence of thousands of freshwater fish species (Katano et al. 2006, Butchart et al. 2010, Lierman et al. 2012, Humphries and Walker 2013, Kroon and Phillips 2015). Importantly, movement within and between river, stream and estuarine environments is essential for fish to access critical habitats for reproduction, food and refuge (Lucas et al. 2009; Humphries and Walker 2013). Anthropogenic modification of waterways can restrict these movements and has led to the fragmentation, decline and local extinction of freshwater fish populations worldwide (Butchart et al. 2010, Humphries and Walker 2013).

Artificial instream barriers such as dams, weirs and culverts, are major contributors to the loss of freshwater biodiversity. Significant work has focused on designing and implementing fish passes for large-scale barriers (dams and weirs) (Loucks 2012, Anderson et al. 2015). However, these large structures are substantially outnumbered by low-head barriers, specifically culverts, that are designed to maintain water connectivity under roads, railways and embankments (Lucas et al. 2009). Recent analysis now estimates that these smaller barriers have a greater cumulative impact on fish populations due to their high abundance within freshwater systems (Januchowski-Hartley et al. 2013). This recognition has fuelled the requirement for remediation strategies that work to improve fish passage through culverts (Duguay and Lacey 2016, Rodgers et al. 2017, Goodrich et al. 2018). Culverts were originally designed to maximise hydraulic capacity with little to no regard for fish passage. Consequently, many culvert designs create a hydraulic barrier to fish movement by increasing water velocities and decreasing surface roughness, creating smoother and more laminar flows through the structure (Rodgers et al. 2014). Recognition of the impact of culverts on fish passage has resulted in the adoption of a range of remediation strategies.

Current culvert remediation strategies include increasing the structure’s cross-sectional area to slow water velocities, addition of baffles and ropes, and roughening of the channel bed with naturalistic or artificial substrates (Table 1)(Chanson and Uys 2016, David et al. 2013, Goodrich et al. 2018, Rodgers et al. 2017, Slawski & Ehlinger 1998). Strategies employing bed roughening and baffles aim to increase the size and frequency of the low velocity boundary layer (BL) at the culvert margins. A boundary layer is formed from the friction between the water and the solid surface over which it flows. Culvert remediation strategies aim to increase the size of the boundary layer, which creates reduced velocity zones (RVZ’s) that fish can exploit (Johnson et al. 2012, Rodgers et al. 2017, Goodrich et al. 2018). These low velocity regions act as energetically favourable movement pathways for fish by reducing the energetic costs associated with swimming in an otherwise high velocity environment. These strategies have proved effective at improving culvert traversability in a number of large bodied commercially important fish species (Johnson et al. 2012), and more recently, for some small-bodied native Australian fishes (Rodgers et al. 2017, Goodrich et al. 2018). In addition to increasing the size of RVZs, baffles, bed roughening and ropes, all create turbulence (Table 1). Turbulence can be defined as the pattern of fluid movement characterised by chaotic changes in flow caused by the interaction of otherwise laminar flow with objects in its path, creating eddies and vortices in its wake. Different fish species respond differently to turbulence, some unfavourably with reduce swimming performance and rates of successful passage (Pavlov et al. 1994, Pavlov et al. 2000, Lupandin 2005, Goodrich et al. 2018). In contrast, some species are able to enhance swimming performance in turbulent flows by utilising eddies (swirling water pockets) to propel themselves forward against water flow via a swimming mode called kármán gaiting (Liao *et al.* 2003a; Liao *et al.* 2003b; Taguchi and Liao 2011). However, for this to strategy to be effective, the eddies must be similar in size to the fish. This restricts the postive effect of turbulance to a certain size class at a particular water velocity, for a specific baffle design. Turbulence does not benefit all fish species, likely due to the interspecific differences in swimming mode, morphology and ecological niche (Goodrich et al. 2018).

**Table 1.** Attributes of current culvert remediation strategies

| Remediation strategy | How it works to increase fish passage | Pros | Cons |
| --- | --- | --- | --- |
| **Increasing structure size** | • Increased cross-section<br>• Reduces velocities through structure | • Reduces risk of upstream flooding | • Increased capital expenditure<br>• Refurbishment of current structure required |
| **Baffles** | • Solid structures generally perpendicular to flow<br>• Provide low velocity rest areas<br>• Create turbulence for kármán gaiting | • Retrofitting opportunities<br>• Cost effective to implement | • Prone to foul with debris<br>• Reduced culvert discharge capacity<br>• Turbulence created is only beneficial for a narrow size class of fish, which is dependent on baffle design and flow characteristics |
| **Bed roughening** | • Created by adding natural rocky substrate<br>• Disrupts laminar flow across an otherwise smooth surface to increase the boundary layer. This creates a low velocity zone near the substrate. Turbulence is also enhanced aiding small-bodied fish to kármán gait | • Can retrofit<br>• Cost effective<br>• Minimal impact on culvert discharge | • Not all fish benefit from turbulence |
| **Addition of ropes** | • Increases surface area roughness<br>• Provides substrate for climbing species | • Can retrofit<br>• Cost effective<br>• Minimal impact on culvert discharge | • Limited number of species |

While altering velocities and turbulence through the various remediation strategies focuses on the biotic requirements, there are complementing concerns about the impacts of remediation strategies on the civil functionality of culverts, that being their ability to discharge water in a cost effective manner. Both baffles and bed roughening compromise culvert functionality by reducing culvert discharge capacity (Olsen & Tullis, 2013), which can have flow-on impacts on upstream flooding. Additionally, both baffles and bed roughening can also increase the likelihood of debris build up and clogging. This increases the maintenance costs associated with culverts remediated via either strategy and can create a physical barrier to fish passage. Taking into account these civil concerns, the positive effect of larger RVZ, and the mixed effects of turbulence, we designed and tested novel remediation strategies to improve fish passage through culverts.

Previously, we identified that Australian small-bodied and juvenile fish predominantly utilise the RVZ associated with culvert corners (Goodrich et al. 2018). Here the boundary layers of the wall and bed merge to create a larger RVZ that is exploited by the fish to reduce their energy expenditure. Subsequently, we developed novel lateral beam designs that run down the length of the channel to increase the size of the RVZ (Fig. 1). We hypothesised that the longitudinal beams would increase the boundary layer effect to increase the size of RVZ and provide low velocity movement paths for fish. It was hypothesised that fish would actively seek out these regions to enhance their swimming endurance and traversability. Two beams (square and rounded) and a ledge design were created, aimed to enhance the size of the RVZ adjacent to the wall of a 12 m flume while minimising excessive turbulence. We compared these to two triangular baffle treatments, which have been shown to have the best compliance with the civil requirements of available baffle designs (Cabonce et al. 2017, Chanson & Uys 2016, Wang & Chanson 2017).

**Figure 1.**
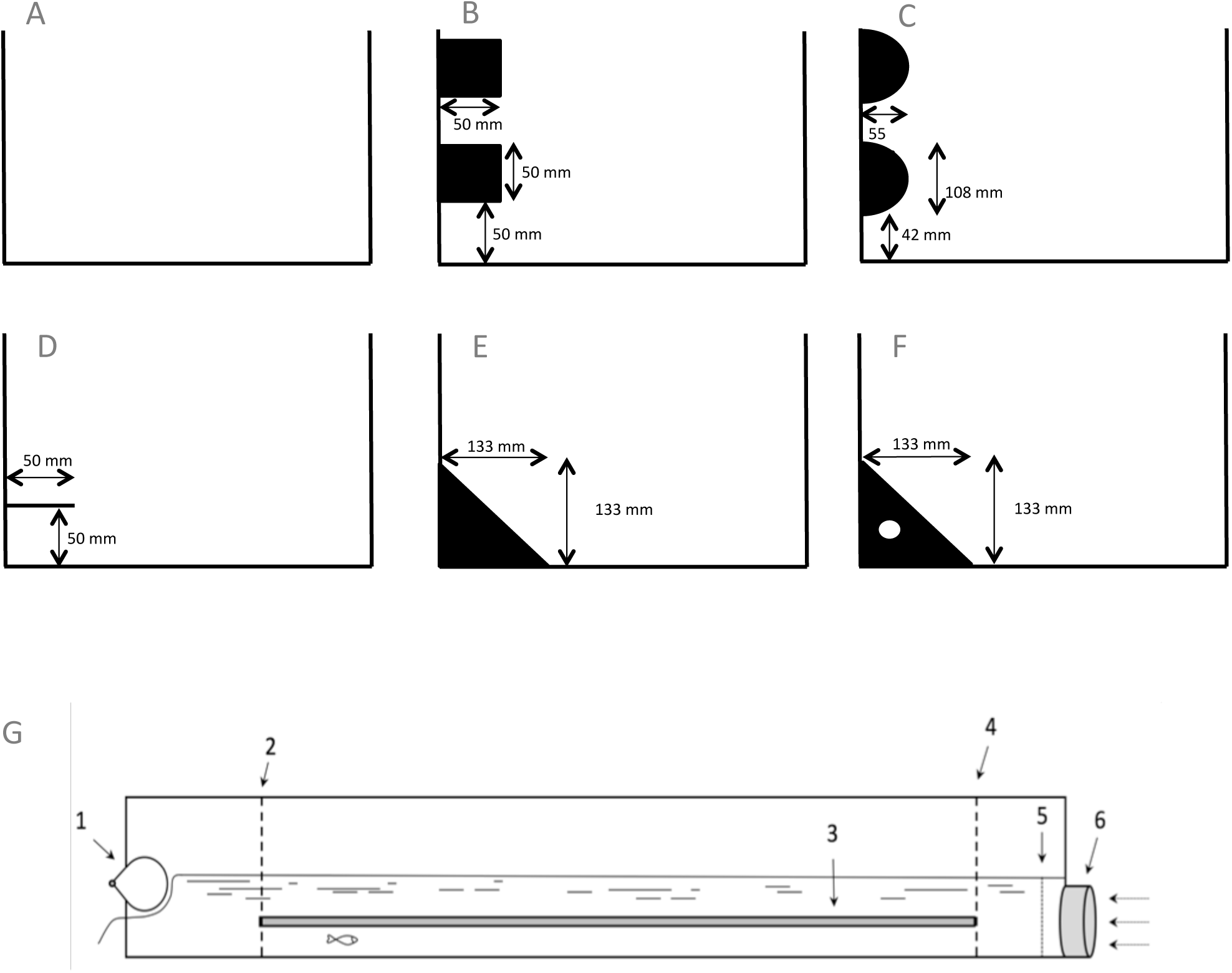
Schematic diagram of flume setup and cross section profiles of the designs tested. We tested a control channel (A) against a square beam (B), circular beam (C), ledge (D), baffle (E) and baffle with hole (F). B, C and D ran longitudinally through the channel, while E and F were spaced 0.66 m apart. Schematic representation of 12 m glass flume (G) used throughout swimming trials showing the depth adjustment gate (1), wire screen to capture fish upon fatigue (2), square beam as example of longitudinal channel modification (3), wire barrier to prevent fish entering inlet chamber (4), flow straighteners (5) and the water inlet (6). Not drawn to scale.

## Methods

### Species choice and husbandry

Fishway research has focused predominately on large bodied, commercially important species like salmon and sturgeon (e.g. Bates and Powers 1998; Taguchi and Liao 2011; Johnson et al. 2012; Lacey et al. 2012, Downie and Kieffer 2017, May and Kieffer, 2017, Erkinaro et al. 2017), with small bodied and juvenile fishes representing an important knowledge gap. Small-bodied fish species represent the most threatened size range of fish (Kalinkat et al. 2017, Olden et al. 2007, Ripple et al. 2017), the loss of which would have lasting impacts on ecosystem functionality, health and services (Mouillot et al. 2013, Rodríguez-Lozano et al. 2015). We chose six small bodied or juvenile (<10 cm total length) native Australian fish species to quantify the effect of our culvert remediation designs: These species are endemic to Australia and represent a variety of different body morphologies, swimming performance, and ecological habits, allowing them to act as proxy species for informed management decisions.

Juvenile *Macquaria ambigua*, (*n* = 90; TL: mean ± s.e. 54.99 ± 0.77 mm; range: 23-76 mm), *Maccullochella peelii* (*n* = 75; TL: mean ± s.e. 73.54 ± 0.78 mm; range: 60-95 mm), *Tandanus tandanus* (*n* = 90; TL: mean ± s.e. 65.23 ± 0.65 mm; range: 45-81 mm) and adult *Hypseleotris compressa* (*n* = 90; TL: mean ± s.e. 59.92 ± 0.90 mm; range: 43-82 mm), *Ambassis agassizii* (*n* = 90; TL: mean ± s.e. 59.29 ± 0.62 mm; range: 40-72 mm) and *Pseudomugil signifer* (*n* = 90; TL: mean ± s.e. 41.81 ± 0.31 mm; range: 35-49 mm) were obtained from commercial hatcheries in southeast Queensland, Australia. Fish were held in 40 L aquariums that formed part of three 1000 L recirculating systems. Water temperature was maintained at 25°C ± 1°C, and fish were exposed to a 12:12 h - light:dark cycle. Fish were fed to satiation daily using commercially sourced fish food pellets (Hikari^™^ micro-wafers) and frozen bloodworms (*Chironomidae)*. All fish were fasted for 24 hours prior to each swimming performance trial to ensure a post-absorptive state (Norin *et al.* 2014).

### Channel remediation designs

Swimming trials were conducted within a 12 m flume (dimensions: 12 × 0.5 × 0.30 m; L × W × H; Fig. 2) which formed part of a 40 000 L recirculating fish swimming facility at The University of Queensland’s Biohydrodynamics Laboratory. Water within the swimming facility was maintained at a constant temperature of 25°C ± 1°C using an industrial heater/chiller (Oasis C58T-Vb, New Zealand). Fish were swum in six treatments including a control channel with PVC sheeting lining the walls (Fig 1). The three novel designs were square beams (12 × 0.5 × 0.5 × m; L × W × H), circular beams (12 × 0.55 m; L × radius) and a ledge (12 ×.05 × 0.05 m; L × W × PVC thickness). Initially we included just the triangular baffle (Isosceles triangle: 0.133 m height). The very poor performance of all species swum in the baffle treatment led to the inclusion of the baffle with hole (Isosceles triangle 0.133 m high, hole diameter 0.013mm, located 4 cm from sidewall and above channel bed). This design was to allow the space before the baffle to ventilate to reduce the strong recirculating turbulence that was disorientated fish. (H. Chanson 2017, personal communication). For both baffle treatments, the baffles were spaced 0.66 m apart. All of the designs were fabricated using polyvinyl chloride (PVC).

**Figure 2.**
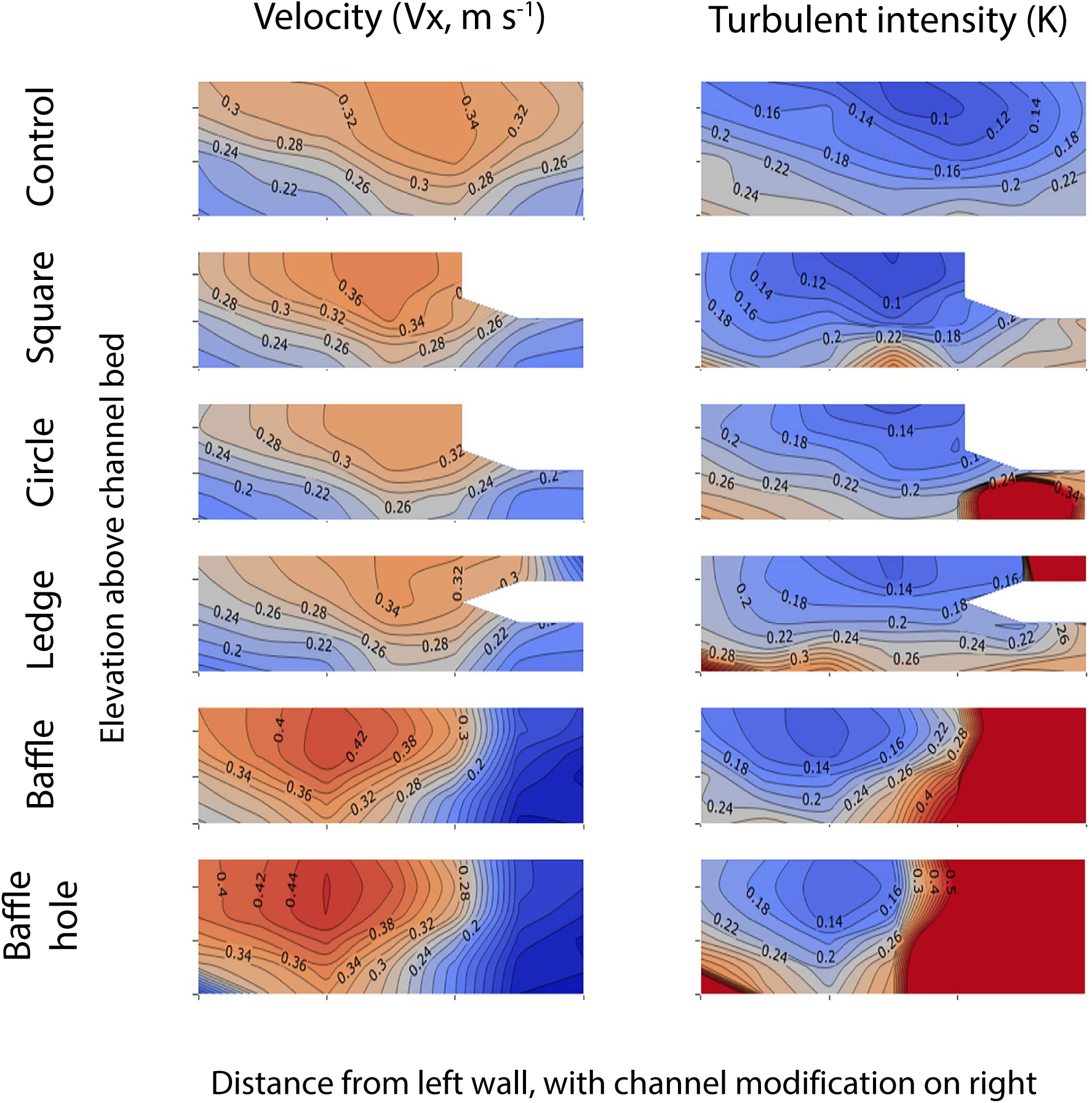
Contour curves of constant longitudinal water velocity (Vx, m s ^−1^) and turbulent intensity (K) in the control channel and remediation treatments (square beam, circle beam, ledge, baffle and baffle with hole). K has no units as it value relative to the mean velocity, for example a 0.2 value represents a 20% fluctuation. Bulk water velocity in the channel was 0.3 m s ^−1^. Modifications were fitted to the right wall of the channel, with white spaces indicating areas that were not accessible to the ADV.

### Swimming performance

The endurance swimming capacity, or time taken to fatigue at a set velocity, was recorded for each individual across each modification treatment and smooth control (*n*=15 per species per modification). All swimming trials lasted for a maximum of 60 min, with trials equal to or greater than 60 mins being treated as censored or unobservable data. No fish were swum in the same treatment more than once, and *M. ambigua, T. tandanus, H. compressa, A. agassizii* and *P. signifer* were given a resting period of at least 14 days between swim trials. Due to rapid growth of juvenile *M. peelii*, individuals were given a resting period of 7 days between swim trials. Each species was swum at bulk water velocities relative to their known Ucrit value, which is the maximum sustained swimming speed a fish is capable of maintaining for 5 min. As swimming fish in different equipment results in different swimming performance values (Kern et al. 2017), we swam the control treatment first to ensure fish fatigued in less than 5 min. Subsequently, *M. ambigua, T. tandanus,* and *H. compressa* were swum at a bulk water velocity of 0.4 m.s^−1^, while *A. agassizii* and *P. signifier* were swum at 0.5 m.s^−1^ and *M. peelii* 0.6 m.s^−1^. These velocities were above the Australian national guidelines for culvert design of 0.3 m.s^−1^ (Fairfull and Witheridge 2003).

### Fish behaviour

Traversable water velocity models can be used to determine the theoretical maximum traversable water velocity that a fish can traverse through culverts of differing lengths (Rodgers et al. 2014, Goodrich et al. 2018). While these models are able to provide recommendations of the maximum water velocity through a culvert of a certain length for a species, they are unable to determine whether the structure will promote fish movement. To give insight into the fish’s behaviour we recorded the traversability of each individual. Traverse success was defined as the ability of the individual to move through 8 m (average length of a culvert within New South Wales, Australia, waterways ranges from 8 – 10 m in length) of the channel without encouragement. Additionally, to determine how the fish were utilising each modification, fish position was also timed during each swimming trial to quantitate if the remediation designs were being used by the fish. Utilisation was defined as the fish swimming underneath, behind, above or directly adjacent to the modification that would potentiate utilising the RVZ created by the design. Percent time spent using the modification was then recorded from the total swim time for each individual.

### Hydraulics

An acoustic Doppler velocimeter (ADV) (Vectrino Laboratory Version, Nortek, serial number: VNO 1686) equipped with a three-dimensional side-looking head was used to determine the boundary layer and turbulent properties of fluid flow in the 12 m channel for each treatment. A 7 x 6 grid (0.5, 3, 12.4, 24.75, 37.1, 46.5 and 49 cm from the left wall and 0.58, 1, 2.5, 5.5, 8.5 and 10 cm depth) of point velocities and turbulent intensity (K) were made at 6 m along the length of the channel. The position of the probe was controlled by a fine adjustment mount connected to a digimatic scale (HAFCO, Taiwan). The ADV was set with a transmit length of 0.3 mm and a sampling volume of 1.5 mm height. The signal was sampled at 200 Hz for 180 s, and a maximum of 36000 samples. ADV data was post processed using WinADV (version 2.030) to remove communication errors, average signal to noise ratio less than 5 dB, and average correlation values < 60%.

Turbulence intensity (K) was calculated using the following equation adapted from Grinval’d and Nikora, (1988), Lupandin (2005), and Goodrich et al (2018):

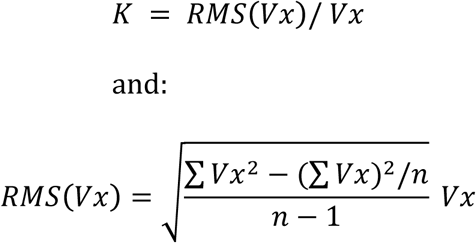

Where K is the turbulent intensity, RMS (Vx) is the root mean square deviation from the time-averaged velocity Vx.

To determine the effect of each modification on the hydrodynamics of fluid flow, heat maps of turbulent intensity and point velocities were created using DPlot graphing software (Version 2.3.5.5, 2017).

### Statistical analyses

All statistical analyses were performed using R version 3.2.3 (R Development Core Team, 2017) in the RStudio environment (Version 1.0.143). The Survival package (Therneau, 2015) was used to analyse endurance data by estimating the probability of fatigue over time for each of the channel treatments. The ZOIB package (Liu and Kong, 2015) was used to analyse the utilisation data using a zero one inflation model. Traversability was analysed using a binomial generalized linear model with traverse success as the response variable and remediation strategy and species as predictors. Statistical significance was defined at *p* < 0.05.

## Results

### Hydraulics

We initially quantified the fluid flow in the control channel and subsequently with each channel modification. Velocity contour plots of channel cross-sections show that the square, circle and ledge designs all altered point velocities and expanded the RVZ in the space adjacent to the wall and channel bed, when compared to the same area in the control channel (Fig. 2). These data also supported our prediction that the square, circle and ledge designs would cause minimal change to the rest of the cross-sectional velocity profile compared to the control channel (Fig. 2). Both baffle treatments were shown to create the largest RVZ along the modified wall, but this corresponded with higher velocities across the rest of the channel cross-section in order to maintain discharge volume. The large change in point velocities observed in the two baffle treatments corresponded with the greatest increases in turbulent intensity (K) (Fig. 2). The turbulent intensities quantified in both baffle treatments were up to two orders of magnitude greater than the control channel (Fig. 2). The turbulence profile of the square modification was most similar to the control channel, with overall increases in turbulent intensities created by both the circle and ledge designs. The circle modification created an area of increased turbulent intensity under the modification but the RVZ associated with the wall was still present. An ANOVA between the point velocities showed no significant differences between treatments, meaning the bulk channel velocity (the average channel velocity) was not altered by any of the designs tested. Comparing the turbulent values between treatments revealed a significant increase caused by the baffles with holes (ANOVA, p > 0.0001) and notable increase cause by baffles (p = 0.06), when compared to the control channel. The comparison of turbulent values associated with the square, circle and ledge, with the control channel supported that the profiles displayed high similarity (ANOVA, all p > 0.98). Overall the square design was able to create the largest RVZ with minimal turbulence generation, by merging the boundary layers of the bed, wall and bottom surface of the beam to create a reduced velocity channel for fish to utilise.

### Fish swimming endurance

The average endurance swimming times (seconds ± SD) in the control channel were 88.2 ± 10.3, 17 ±1.4, 63 ± 6.7, 50.7 ± 7.4, 47.8 ± 11.6 and 50.5 ± 3.6 for *Ambassia agassizii, Pseudomugil signifier, Hypseleotris compressa, Macquaria ambigua, Tandanus tandanus* and *Maccullochella peelii* respectively (Fig. 3). Total body length had a significant effect on swimming endurance in *P. signifier* (*X*^2^ = 9.24, *p* = 0.002) but had no effect on swimming endurance for any other species (*A.agassizi*: *X*^2^ _*7*_= 2.22, *p* = 0. 14; *H. compressa*: *X*^2^ _*7*_= 0.58, *p* = 0.44; *M. ambigua*: *X*^2^ _*7*_= 0.30, *p* = 0.60; *T. tandanus*: *X*^2^ _*7*_= 0.70, *p* = 0.4; *M. peelii*: *X*^2^ _*6*_= 1.90, *p* = 0.17). Body mass had a significant effect on the swimming endurance of *M. peelii* (*X*^2^ _*6*_= 9.48, *p* = 0.002) and *P. signifier* (*X*^2^ _*7*_= 4.47, *p* = 0.03), but had no effect on swimming endurance performance for any other species (*A. agassizi*: *X*^2^ _*7*_= 1.96, *p* = 0.16; *M. ambigua*: *X*^2^ _*7*_= 1.1, *p* = 0.3; *T. tandanus*: *X*^2^ _*7*_ = 1.78, *p* = 0.2; *H. compressa*: *X*^2^ _*7*_= 0.88, *p* = 0.35).

**Figure 3.**
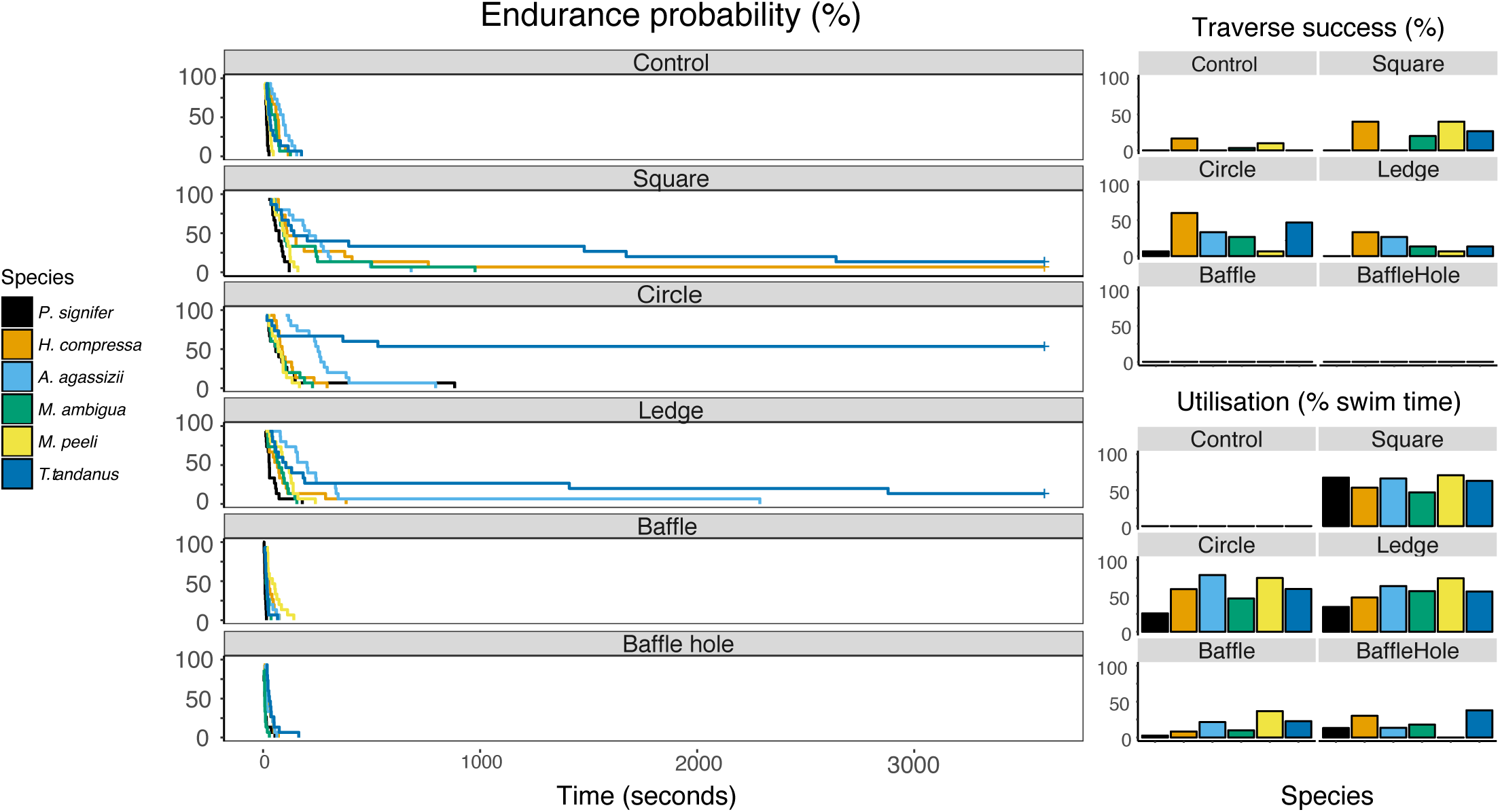
Endurance probability of the six species swum under each test condition (+ sign indicates some fish of that species swam for the maximum time of 3600 s.in that design and the data was censored accordingly). All of the beam style designs showed longer endurance times when compared to the control channel. Overall both baffle designs had a negative effect on endurance times. This pattern was repeated with the rates of traverse success (proportion ± s.e.), which represents the number of fish that were able to successfully swim 8 m of the test channel. Average time spent utilising channel modifications (proportion (%) ± s.e.), with utilisation defined as the fish being positioned adjacent to the modification and within the RVZ. Utilisation time was not recorded for the control channel.

Relative to the smooth control channel, the semi-circle design significantly increased the swimming endurance performance of *A. agassizii* (*X*^2^ _*7*_= 21.16, *p* = <0.001), *P. signifier* (*X*^2^ _*7*_= 46.76, *p* = < 0.001), *M. ambigua* (*X*^2^ _*7*_= 5.06, *p* = 0.02), *T. tandanus* (*X*^2^ _*7*_= 62.03, *p* = <0.001) and *M. peelii* (*X*^2^ _*6*_ = 28.4, *p* = < 0.001)(Fig. 3). This design was most beneficial to *T. tandanus* and *P. signifier* with the introduction of the beam increasing their mean swimming endurance performance above that of any other modification when compared to the control channel.

The longitudinal ledge significantly increased the swimming performance, above that in the control condition, of four of the six examined species: *A. agassizii* (*X*^2^ _*7*_= 39.82, *p* = < 0.001), *P. signifier*(*X*^2^ _*7*_= 13.46, *p* = < 0.001), *T. tandanus* (*X*^2^ _*7*_= 31.14, *p* = < 0.001), and *M. peelii* (*X*^2^ _*7*_= 79.21, *p* = < 0.001)(Fig. 3). This modification was most beneficial to *M. peelii* and *A. agassizii* with the introduction of the ledge increasing their mean swimming endurance performance above that of any other modification when compared to the control channel.

The square longitudinal beam was the only modification to significantly increase the swimming endurance performance of all species when compared to the control channel (*A. agassizii* (*X*^2^ _*7*_=16.08, *p* = < 0.001), *P. signifier* (*X*^2^ _*7*_= 27.55, *p* = < 0.001), *T. tandanus* (*X*^2^ _*7*_= 37.74, *p* = < 0.001), *M. peelii* (*X*^2^ _*6*_= 21.90, *p* = < 0.001), *M. ambigua* (*X*^2^ _*7*_= 40.92, *p* = < 0.001), *H. compressa* (*X*^2^ _*7*_= 31.19, *p* = < 0.001)(Fig. 3). This modification was most beneficial to *H. compressa* and *M. ambigua* with the introduction of the beam increasing their mean swimming endurance performance above that of any other modification when compared to the smooth control channel.

Both baffles designs were the only modifications used throughout this study that did not provide a performance benefit to any species, and even decreased the average endurance times of some species when compared to the smooth control channel (Fig. 3). The baffles significantly decreased the swimming endurance of *A. agassizii* (*X*^2^ _*7*_= 20.61, *p* = < 0.001), *P. signifier* (*X*^2^_7_= 6.02, *p* = 0.01), *H. compressa* (*X*^2^ _*7*_= 5.87, *p* = 0.02), *M. ambigua* (*X*^2^ _*7*_= 23.60, *p* = < 0.001) and *T. tandanus* (*X*^2^ _*7*_= 3.94, *p* = 0.047). Likewise, the baffles with holes significantly decreased the swimming endurance of *A. agassizii* (*X*^2^ _*7*_= 21.72, *p* = < 0.001), *H. compressa* (*X*^2^ _*7*_= 7.09, *p* = 0.007), *M. ambigua* (*X*^2^ _*7*_= 48.34, *p* = < 0.001), and *T. tandanus* (*X*^2^ _*7*_= 4.25, *p* = 0.04 when compared to thecontrol channel.

### Utilisation of channel modifications by fish

Upon beginning a swim trial, individual fish would generally explore the channel bed. Once the RVZ was discovered, most fish would utilise it until fatigue. Utilisation was defined by the body position of the fish being adjacent to a surface where it was in a RVZ created by either the beam or baffle designs. Most species spent similar times utilising the longitudinal remediation strategies and less time utilising either baffle designs (Fig. 3). Specific comparisons between each of the channel modifications revealed that *A. agassizii, H. compressa, M. ambigua, T. tandanus* and *M. peelii* spent significantly more time utilising the square beam when compared to the baffle (p < 0.0001; p < 0.0001; p = 0.04; p = 0; p = 0.08 respectively). Likewise, *A. agassizii, P. signifier, H. compressa, T. tandanus* and *P. signifier* spent significantly more time utilising the square beam when compared to baffles with holes (p < 0.0001; p = 0.04; p = 0.01; p = 0.008; p = 0.04 respectively).

The circular modification also showed significantly higher utilisation by several species when compared to the baffle and baffle with hole. *A. agassizii, H. compressa* and *T. tandanus* spent significantly more time using the circle than the baffle (all p < 0.0001) and baffle with hole (all p < 0.0001). Furthermore, *M. ambigua*, and *M. peelii* spent significantly more time utilising the circle modification when compared to the baffle (p < 0.0001; p < 0.0001; p = 0.01 respectively).

Between the two baffle designs, *M. ambigua* was the only species with a significant difference in utilisation times, spending significantly more time using the baffles with holes, than the baffles (p = 0.02). There was no significant difference between the average time spent utilising the square and circle, and square and ledge modifications, for all species. *H. compressa* was the only species to spend significantly more time using the circular modification when compared to the ledge (p = 0.04).

Comparing the ledge with the baffle designs showed that *H. compressa, T. tandanus, M. ambigua* and *P. signifier* all spent significantly more time using the ledge (all comparisons p < 0.0001).

### Traversability

Averaging over the species swum, the square (p = 0.001), semicircle (p < 0.0001) and ledge (p = 0.047) all significantly improved the traverse success rates when compared to the smooth control channel. There was no significant difference between the beam designs when averaged across species. The circular design significantly improved the traverse success of *P. signifier, H. compressa, A. agassizii, M. ambigua* and *T. tandanus* (all p < 0.0001). Although *M. peeli* showed a significant decrease in traverse success in the circular design compared to the smooth control channel (p < 0.0001). The square design showed a significant increase in traverse success for *P. signifier, M. peeli, A. agassizii, M. ambigua* and *T. tandanus (*all p < 0.038). No fish were able to traverse the test channel in either of the baffle designs.

### Scaling of the RVZ under the square design with bulk channel flow

Further hydrological characterisation was done on the square modification to determine how the RVZ under the beam responded to changes in bulk channel velocity relative to the same corner in the smooth control channel (no beams or baffles). The bottom surface of the square modification caused the boundary layers associated with the wall and bed to merge and created a pocket of reduced velocity (Fig. 4). This reduction was exacerbated with decreasing bulk channel velocity (Fig. 4), indicating that greater benefit to fish is produced at lower bulk channel velocities. Over the range of bulk channel velocities measured here, the mean Vx under the beam was approximately 20 % lower than in the main channel body.

**Figure 4.**
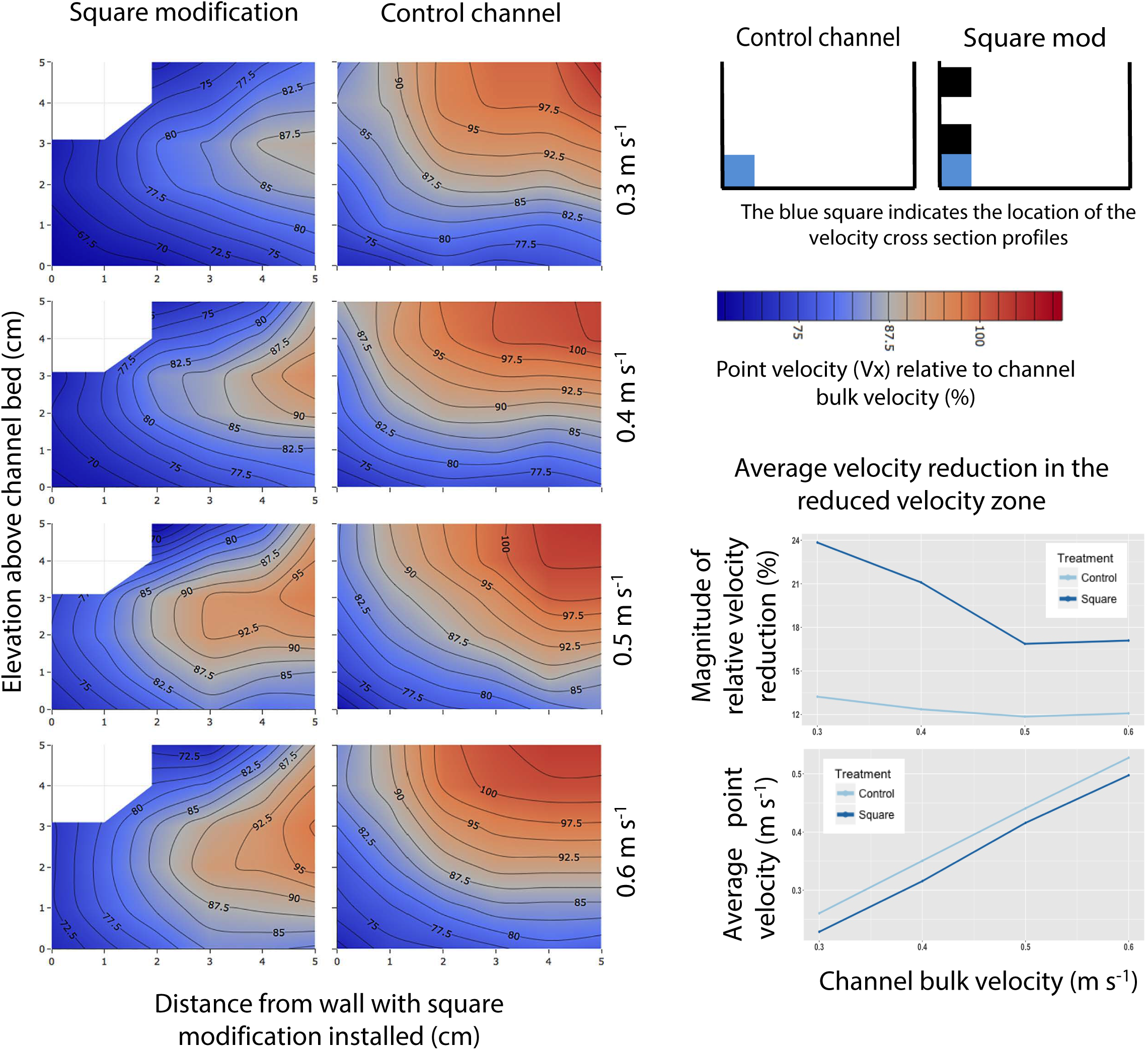
Scaling of the RVZ under the square modification with increasing bulk channel velocity. Values are presented at percent reduction of bulk channel flow.

## Discussion

The novel beam designs all provide a significant improvement in the endurance and traversability of a wide range of small-bodied and juvenile fish species when compared to either baffle designs or the control channel. Further, we show that baffles can have a negative impact on small fish, largely due to the generation of excessive turbulence at high bulk velocities relative to their swimming capacity. By creating a reduced velocity channel with the longitudinal beam, we avoid the issues created by excessive turbulence and increase the maximum bulk channel flow that any fish can swim against, regardless of morphology and ecology.

The longitudinal beam designs generated an expanded low velocity zone between the beam and the base of the channel that reduced the velocity of water by as much as 33 % of the bulk channel flow. The majority of fish species showed a clear preference for, and performance improvement in the presence of, the longitudinal beams designs compared to both baffle designs. This was because the baffle designs generated considerable turbulence at high bulk channel velocities that had a negative impact on small fish. Although it could be argued that baffles may still provide benefits for fish at slower bulk channel velocities where turbulence is reduced, low velocity flows are less likely to be problematic for fish. Additionally, the magnitude of the velocity reduction provided in the RVZ increases with reduced bulk channel flow. Simply, the slower the overall channel flow, the more effective the square beam becomes at providing a RVZ for fish to utilise. Given this, the square beam design provides an alternative to baffles, which may be unsuitable for promoting the passage of some small bodied or juvenile fish through culverts where high discharges occur. By contrast, all longitudinal beam designs produced minimal turbulence and had little impact on channel discharge capacity indicating that they may prove more effective for remediating culverts to improve the upstream passage of small fish whilst permitting relatively high bulk velocities.

Irrespective of differences in body size, shape, swimming capacity and mode, all tested fish species were able to derive a performance benefit at high water velocities in the presence of the square beam design. In contrast, baffles impaired fish swimming performance at high bulk velocities likely due to the high degree of turbulence generated by water flowing over and around the baffles. Excessive turbulence creates hydraulic barriers to fish movement and increases energetic cost of swimming (Enders et al. 2011, Yuan et al. 2017) unless the size of the eddies are comparable to the fish body size (Pearson 2006). As eddy size will change with channel bulk velocity (Baki et al. 2015), and natural streams contain complex assemblages of species and size classes (Dudgeon et al. 2005, Winemiller and Leslie 1992), the ability of baffles to provide a positive effect is limited to a small bulk velocity for an individual fish. Additionally, the size of, and spacing between baffles determines the degree of turbulence generated and must be optimized for target fish species and size class (Feurich et al. 2012, Rajaratnam et al. 1988), adding complexity into the assessment of their benefit. This limits their efficacy for promoting fish passage for small-bodied fish. In contrast, the novel beam designs appear to avoid these issues by creating a consistent reduced velocity channel with minimal generation of turbulence along the full length of the channel. Specific to our study, this resulted in an increase in endurance and traversability for all six fish species in most of the beam treatments, and in a broad context, could greatly increase the maximum bulk channel velocity under which any small fish could successfully traverse a culvert *in situ*.

The quantification of fluid flow characteristics showed that, as predicted, all longitudinal beam designs increased the size of the RVZ when compared to the smooth culvert control channel. These RVZ’s form due to the friction of the water flowing against a surface and increase in size when two surfaces run adjacent to one another and their respective boundary layers merge (Goodrich et al. 2018). The use of RVZ’s by fish is well supported here and in the literature (Goodrich et al. 2018, Powers et al. 1997, Johnson et al. 2012, Rodgers et al. 2017). The RVZ provides an energetically favourable movement path for fish by reducing the energetic cost associated with upstream swimming, increasing their endurance capacity (Johnson et al. 2012). An increase in swimming endurance was demonstrated for at least one species in all of the beam style designs, but notably all species benefited from the square design. By contrast, neither baffle design was able to improve fish swimming endurance capacity, suggesting that at high water velocities, these baffles may create levels of turbulence which exceed the swimming capabilities of some small bodied fish species.

In contrast to the triangular baffles, fish were very often found to be within or to associate closely with, the beam designs. We propose that in addition to the favourable energetic conditions created by these beam designs, the covered spaces under the modifications may act as behavioural refuge for small fish. It is known that habitat structure influences prey mortality (Klecka et al. 2014), and fish that seek refuge from predators are less likely to be captured and consumed (Savino and Stein 1982, Denno et al. 2005). We suggest that a combination of energetic benefits and behavioural responses contributed to the amount of time each species spent utilising the different modifications tested throughout this study.

The longitudinal beam designs provided a reduced velocity zone in relatively high water flows that was between 16 and 30% lower than that of the main channel flow. Although the relative magnitude of the RVZ reduction decreased with increasing bulk velocity, that they still facilitated fish passage at high water flows shows that they can effectively provide a velocity refuge for fish that can be exploited for upstream passage. Moreover, all of the longitudinal bean designs tested here had negligible impact on channel discharge capacity and so are likely to conform to civil requirements and performance capacity. The longitudinal beams designs are also less likely to trap debris like baffles do, which may improve their appeal to infrastructure managers and increase their utility for fish passage in new and remediated culverts.

Despite the negative findings reported here for small bodied fish, baffles of various designs remain one of the most widely implemented culvert remediation strategies (Kaptizke 2010). This is due to the positive effect baffles have on improving the passage of large, commercially important fish species, with the literature heavily weighted towards salmonids (Forty et al. 2016). Large bodied fish would be unlikely to benefit from the beam designs tested here as increasing the size of the void beneath the beam would likely reduce the relative change in water velocity. Moreover, the water flow velocities used in the current study would be unlikely to impact the passage of large bodied fish, assuming water depth was sufficient. However, we have to date, only tested one spacing configuration in relation to fish passage. While effective for the majority of small fish examined here, combinations of remediation designs, dependent on site-specific community structure, may be appropriate to ensure all commercially and/or ecologically species can traverse culverts at high bulk flows.

The square, circle and ledge remediation designs tested here all improved the ability of small fish to swim in a culvert like channel against high bulk velocities, while having negligible impact on the civil requirements and performance capacity. The square longitudinal ledge was the only treatment to significantly increase fish swimming endurance across all species when compared to the smooth control channel, and significantly increased traversability in four of six species. Of the designs tested, we recommend that the square beam modification be chosen for further optimisation of the RVZ to maximise the size and velocity reduction relative to the bulk channel flow. Furthermore, we hypothesise that adding minor levels of texture within the cavity of the beam design will increase the effect of the boundary layers merging, and achieve a greater relative reduction in velocity in the RVZ. This simple design is likely to be cost effective to retrofit existing structures, will likely require minimal maintenance costs once inserted, and based on our preliminary work, should have minimal impact on culvert discharge capacity. The application of this concept could be extended from box culverts, to any structure where two smooth surfaces intersect, for example within fishways such as vertical slots.

## Acknowledgements

Funding was by an Australian Research Council Linkage Grant [LP140100225] to CEF and MG and conducted under Australian Ethics Committee (AEC) approval number SBS/312/15/ARC. The authors would like to thank Dr Simon Blomberg for statistical advice. We appreciated discussions with Prof. Hubert Chanson. We declare no conflicts of interest.

## References

Anderson, J., Faulds, P., Burton, K., Koehler, M., Atlas, W. and Quinn, T. (2015). Dispersal and productivity of Chinook (*Oncorhynchus tshawytscha*) and coho (*Oncorhynchus kisutch*) salmon colonizing newly accessible habitat. Canadian Journal of Fisheries and Aquatic Sciences, 72(3):454–465.

Baki, A. B. M, Zhu, D. Z. and Rajaratnam, N. (2015). Turbulence characterisitics in a rock-ramp type fish pass. Journal of Hydraulic Engineering, 141(2):04014075

Butchart, S. H. M., Walpole, M., Collen, B., van Strien, A., Scharlemann, J. P. W., Almond, R. E. A., Baillie, J. E. M., Bomhard, B., Brown, C., Bruno, J., Carpenter, K. E., Carr, G. M., Chanson, J., Chenery, A. M., Csirke, J., Davidson, N. C., Dentener, F., Foster, M., Galli, A., Galloway, J. N., Genovesi, P., Gregory, R. D., Hockings, M., Kapos, V., Lamarque, J.-F., Leverington, F., Loh, J., McGeoch, M. A., McRae, L., Minasyan, A., Hernández Morcillo, M., Oldfield, T. E. E., Pauly, D., Quader, S., Revenga, C., Sauer, J. R., Skolnik, B., Spear, D., Stanwell-Smith, D., Stuart, S. N., Symes, A., Tierney, M., Tyrrell, T. D., Vié, J.-C. and Watson, R. (2010). Global biodiversity: indicators of recent declines. Science 328:1164–1168.

Cabonce, J., Fernando, R., Wang, H. and Chanson, H. (2017). Using Triangular Baffles to Facilitate Upstream Fish Passage in Box Culverts: Physical Modelling, School of Engineering Report CH107/17, The University of Queensland.

Carpenter, S. R., Stanley, E. H. and Vander Zanden, M. J. (2011). State of the World’s Freshwater Ecosystems: Physical, Chemical, and Biological Changes. Annual Review of Environment and Resources, 36(1):75–99. http://doi.org/10.1146/annurev-environ-021810-094524

Chanson, H. and Uys, W. (2016). Baffle Designs to Facilitate Fish Passage in Box Culverts: A Preliminary Study, ^th International Symposium on Hydraulic Structures, Hydraulic Structures and Water System Management, 1–10. http://doi.org/10.15142/T300628160828

David, B. O., Tonkin, J. D., Taipeti, K. W. T. and Hokianga, H. T. (2013). Learning the ropes: mussel spat ropes improve fish and shrimp passage through culverts. Journal of Applied Ecology, 51(1):214–223. http://doi.org/10.1111/1365-2664.12178

Denno R. F., Finke D. L. and Langellotto G. A. (2005) Direct and indirect effects of vegetation structure and habitat complexity on predator–prey and predator–predator interactions. In: Barbosa P, Castellanos I (eds) Ecology of predator–prey interactions. Oxford University Press, Oxford, UK, pp 211–239

Dudgeon, D., Arthington, A. H., Gessner, M. O., Kawabata, Z., Knowler, D. J., Leveque, C., Naiman, R. J., Prieur-Richard, A., Soto, D., Stiassny, M. L. J. and Sullivan, C. A. (2005), Freshwater biodiversity:importance, threats, status and conservation challenges. Biological Reviews, 81:163–182, https://doi.org/10.1017/S1464793105006950

Duguay, J. M. and Lacey, R. W. J. (2016) Numerical study of an innovative fish ladder design for perched culverts. Canadian Journal of Civil Engineering 43:173–181.

Fairfull, S. and Witheridge, G. (2003), Why do fish need to cross the road? Fish Passage requirements for waterway crossings. NSW DPI, Cronulla.

Farrell, A. P., 2011. Encyclopedia of fish physiology: from genome to environment. Academic Press.

Feurich, R., Boubee, J. and Olsen, N. R. B. (2012). Improvement of fish passage in culverts using CFD, Ecological Engineering, 47:1–8. http://dx.doi.org/10.1016/j.ecoleng.2012.06.013

Forty, M., Spees, J., & Lucas, M. C. (2016). Not just for adults! Evaluating the performance of multiple fish passage designs at low-head barriers for the upstream movement of juvenile and adult trout Salmo trutta. Ecological Engineering, 94:214–224. http://doi.org/10.1016/j.ecoleng.2016.05.048

Goodrich, H. R., Watson, J. R., Cramp, R. L., Gordos, M. A., & Franklin, C. E. (2018). Making culverts great again. Efficacy of a common culvert remediation strategy across sympatric fish species. Ecological Engineering, 116:143–153. http://doi.org/10.1016/j.ecoleng.2018.03.006

Gordon, T. A. C., Harding, H. R., Clever, F. K., Davidson, I. K., Davison, W., Montgomery, D. W., et al. (2018). Fishes in a changing world: learning from the past to promote sustainability of fish populations. Journal of Fish Biology, 92(3):804–827. http://doi.org/10.1111/jfb.13546

Hoerner, S. F. (1965) Fluid dynamic drag. Published by the author, Bakersfield, CA.

Humphries, P. O. and Walker, K. F. (Eds) (2013). ‘Ecology of Australian Freshwater Fishes.’ (CSIRO: Melbourne.)

Januchowski-Hartley S. R., McIntyre P. B., Diebel M., Doran P. J., Infante D. M., Joseph C. and Allan D. J. (2013) Restoring aquatic ecosystem connectivity requires expanding inventories of both dams and road crossings. Frontiers in Ecology and the Environment, 11:211–217.

Johnson E. G., Pearson W. H., Southard S. L. and Mueller R. P. (2012) Upstream movement of juvenile coho salmon in relation to environmental conditions in a culvert test bed. Transactions of the American Fisheries Society, 141:1520–1531.

Kalinkat, G., Jähnig, S. C. and Jeschke, J. M., (2017), Exceptional body size–extinction risk relations shed new light on the freshwater biodiversity crisis. Proceedings of the National Academy of Sciences, https://doi.org/10.1073/pnas.1717087114.

Kapitzke, R. (2010), Fish Passage Planning and Design. Culvert Fishway Planning and Design Guidelines: Part F – Baffle Fishways for Box Culverts. James Cook University, School of Engineering and Physical Sciences.

Katano, O., Nakamura, T., Abe, S., Yamamoto, S. and Baba, Y. (2006), Comparison of fish communities between above- and below-dam sections of small streams; barrier effect to diadromous fishes. Journal of Fish Biology, 68:767–782.

Kern, P., Cramp, R. L., Gordos, M. A., Watson, J. R. and Franklin, C. E. (2017). Measuring Ucrit and endurance: equipment choice influences estimates of fish swimming performance. Journal of Fish Biology, 178, 465–11. http://doi.org/10.1111/jfb.13514

Kroon, F. J. (2005). Behavioural avoidance of acidified water by juveniles of four commercial fish and prawn species with migratory life stages. Marine Ecology Progress Series, 285:193–204.

Kroon Frederieke J. and Phillips Seonaid (2015) Identification of human-made physical barriers to fish passage in the Wet Tropics region, Australia. Marine and Freshwater Research, 67:677–681.

Liao, J. C., Beal, D. N., Lauder, G.V. and Triantafyllou, M. S. (2003a) The Kármán gait; novel kinematics of rainbow trout swimming in a vortex street. Journal of Experimental Biology, 206:1059–1073.

Liao, J. C., Beal, D. N., Lauder, G.V. and Triantafyllou, M.S. (2003b) Fish exploiting vortices decrease muscle activity. Science, 302:1566–1569.

Liao, J. C. and Cotel, A. (2013) Effects of turbulence on fish swimming in aquaculture. Pages 109-127 in P. A. Palstra and V. J. Planas, editors. Swimming Physiology of Fish: Towards Using Exercise to Farm a Fit Fish in Sustainable Aquaculture. Springer Berlin Heidelberg, Berlin, Heidelberg.

Liermann, C. R., Nilsson, C., Robertson, J. and Ng, R. Y. (2012). Implications of dam obstruction for global freshwater fish diversity. BioScience, 62:539–548.

Liu, F. and Kong, Y. (2015). ZOIB: an R Package for Bayesian Inferences in Beta and Zero One Inflated Beta Regression Models, The R Journal, 7(2):34–51

Lovett, R. A. (2014). Rivers on the run. Nature, 511:521–523.

Loucks, E. D. (2012). Providing Fish Passage at the Middle Fork Nooksack River Diversion Dam. World Environmental and Water Resources Congress 2012: Crossing Boundaries, 1546–1558.

Lucas, M. C., Bubb, D. H., Jang, M., Ha, K. and Masters, J. E. G. (2009). Availability of and access to critical habitats in regulated rivers: Effects of low-head barriers on threatened lampreys. Freshwater Biology, 54:621–634.

Lupandin, A.I. (2005) Effect of flow turbulence on swimming speed of fish. Biological Bulletin 32, 461.

May, L. and Kieffer, J.D. (2017) The effect of substratum type on aspects of swimming performance and behaviour in shortnose sturgeon Acipenser brevirostrum. Journal of Fish Biology 90:185–200, https://doi.org/10.1111/jfb.13159

Mouillot, D., Bellwood, D. R., Baraloto, C., Chave, J., Galzin, R., Harmelin-Vivien, M., Kulbicki, M., Lavergne, S., Lavorel, S., Mouquet, N. and Paine, C.T. (2013) Rare species support vulnerable functions in high-diversity ecosystems. PLoS biology. https://doi.org/10.1371/journal.pbio.1001569

Morris, S. A., Pollard, D. A., Gehrke, P. C. and Pogonoski, J. J. (2001). Threatened and potentially threatened freshwater fishes of coastal New South Wales and the Murray-Darling Basin. Report to Fisheries Action Program and World Wide Fund for Nature 33.

New South Wales, Roads and Maritime Services. (2009) Rehabilitation Guideline for Corrugated Steel Culverts. Roads and Traffic Authority, NSW, Australia.

Norin, T., Malte, H. and Clark, T. D. (2014) Aerobic scope does not predict the performance of a tropical eurythermal fish at elevated temperatures. Journal of Experimental Biology 217:244–251.

Norman, J. R., Hagler, M. M., Freeman, M. C. and Freeman, B. J. (2009) Application of a multistate model to estimate culvert effects on movement of small fishes. Transactions of the American Fisheries Society, 138:826–838.

Obregón, C., Lyndon, A. R., Barker, J., Christiansen, H., Godley, B. J., Kurland, S., Piccolo, J. J., Potts, R., Short, R., Tebb, A. and Mariani, S. (2018). Valuing and understanding fish populations in the Anthropocene: key questions to address. Journal of Fish Biology, 92(3), 828–845. http://doi.org/10.1111/jfb.13536

Olden, J. D., Hogan, Z. S. and Zanden, M. J. V. (2007), Small fish, big fish, red fish, blue fish: size-biased extinction risk of the world’s freshwater and marine fishes. Global Ecology and Biogeography, 16:694–701.

Olsen, A. H. and Tullis, B. P. (2013). Laboratory Study of Fish Passage and Discharge Capacity in Slip-Lined, Baffled Culverts. Journal of Hydraulic Engineering, 139(4), 424–432. http://doi.org/10.1061/(ASCE)HY.1943-7900.0000697

Pavlov, D. S., Lupandin, A. I. and Skorobogatov, M. A. (1994) The effect of turbulence degree on the critical current speed of the Gudgeon. Proceedings of the Russian Academy of Sciences, 336:138–141.

Pavlov, D. S., Lupandin, A. I. and Skorobogatov, M. A. (2000) The effects of flow turbulence on the behavior and distribution of fish. Journal of Ichthyology, 40:S232–S261.

Pearson W. H., Southard S. L., May C. W., Skalski J. R., Townsend R. L., Horner-Devine A. R., Thurman D. R., Hotchkiss R. H., Morrsion R. R., Richmond M. C. and Deng, D. (2006) Research on the upstream passage of juvenile salmon through culverts: retrofit baffles. Final Report Prepared for the Washington State Department of Transportation: WSDOT agreement No. GCA2677. Battelle Memorial Institute, Washington.

Rajaratnam, N., Katopdis, C. and Lodewyk S. (1988). Hydraulics of offset baffle culvert fishways, Canadian Journal of Civil Engineering, 15:1043–1051

Ripple, W. J., Wolf, C., Newsome, T. M., Hoffmann, M., Wirsing, A. J. and McCauley, D. J. (2017). Extinction risk is most acute for the world’s largest and smallest vertebrates. Proceedings of the National Academy of Sciences, 114(40):10678–10683, doi.org/10.1073/pnas.1702078114

Rodgers, E. M., Cramp, R. L., Gordos, M., Weier, A., Fairfall, S., Riches, M. and Franklin, C. E. (2014) Facilitating upstream passage of small-bodied fishes: linking the thermal dependence of swimming ability to culvert design. Marine and Freshwater Research, 65:710–719, doi.org/10.1071/MF13170

Rodgers, E. M., Heaslip, B. M., Cramp, R. L., Riches, M., Gordos, M. A., and Franklin, C. E. (2017). Substrate roughening improves swimming performance in two small-bodied riverine fishes: implications for culvert remediation and design. Conservation Physiology, 5(1):1–10. http://doi.org/10.1093/conphys/cox034

Rodríguez-Lozano, P., Verkaik, I., Rieradevall, M., and Prat, N. (2015). Small but Powerful: Top Predator Local Extinction Affects Ecosystem Structure and Function in an Intermittent Stream. PLoS ONE, 10(2), e0117630–16. http://doi.org/10.1371/journal.pone.0117630

Savino J. F. and Stein R. A. (1982) Predator–prey interactions between largemouth bass and bluegills as influenced by simulated vegetation. Transactions of the American Fisheries Society, 111:255–266, doi:10.1577/1548-8659(1982)111<255:PIBLBA>2.0.CO;2

Silva, A. T., Lucas, M. C., Castro-Santos, T., Katopodis, C., Baumgartner, L. J., Thiem, J. D., Aarestrup, K., Pompeu, P. S., O’Brien, G. C., Braun, D. C. Burnett, N. J., Zhu, D. Z., Fjeldstad, H., Forseth, T., Rajaratnam, N., Williams, J. G. and Cooke, S. J. (2017). The future of fish passage science, engineering, and practice. Fish and Fisheries, 19(2):340–362, doi.org/10.1111/faf.12258

Slawski, T. M. and Ehlinger, T. J. (1998). Fish Habitat Improvement in Box Culverts: Management in the Dark? North American Journal of Fisheries Management, 18(18):676–685. http://doi.org/10.1577/1548-8675(1998)018<0676:fhiibc>2.0.co;2

Taguchi, M. and Liao, J. C. (2011). Rainbow trout consume less oxygen in turbulence: the energetics of swimming behaviors at different speeds, Journal of Experimental Biology, 214:1428–1436, doi:10.1242/jeb.052027

Therneau, T. (2015). A Package for Survival Analysis in S. version 2.38, https://CRAN.R-project.org/package=survival.

Vowles, A. S., Anderson, J. J., Gessel, M. H., Williams, J. G. and Kemp, P. S. (2014). Effects of avoidance behaviour on downstream fish passage through areas of accelerating flow when light and dark. Animal Behaviour, 92:101–109, https://doi.org/10.1016/j.anbehav.2014.03.006

Winemiller, K. O. and Leslie, M. A. (1992). Fish assemblages across a complex tropical freshwater/marine ecotone, Environmental Biology of Fishes, 34:29–50

Yuan, X., Jiang, Q., Meng, X., Tu, Z., Zhou, Y. and Huang, Y. (2017). Effect of vertical slit turbulence on metabolism and swimming behavior of juvenile grass carp (ctenopharyngodon idella), Biorxiv, 1–23. http://doi.org/10.1101/190272

